# Hydroxycarboxylic acid receptor 2 (HCAR2/GPR109A) expression and signaling promotes the maintenance of an immunoinhibitory retinal environment

**DOI:** 10.1101/2021.03.15.435493

**Authors:** Folami L. Powell, Amany Tawfik, Pachiappan Arjunan, Deeksha Gambhir Chopra, Mohamed Al-Shabrawey, Nagendra Singh, Ravirajsinh Jadeja, Matthew Kaufman, Malita Jones, Ollya Fromal, Alan Saul, Wan Jin Jahng, Manuela Bartoli, Pamela M. Martin

## Abstract

**Background:** Excessive oxidative stress and related chronic, sub-clinical inflammation is linked causally to the development and progression of degenerative diseases of the retina including diabetic retinopathy, age-related macular degeneration and glaucoma, leading causes of blindness worldwide. The above responses may be related directly to dysregulated retinal immunity and are potentiated by the combined actions of native retinal cells (e.g., retinal pigment epithelial (RPE) and microglial cells) and immune cells infiltrating from the periphery. Maintaining tight regulation of these cells such that effective control of pathogens is accomplished yet uncontrolled inflammation and consequent tissue damage is prevented is extremely important. However, the molecular mechanisms that control this delicate balance are poorly understood. We hypothesize that the hydroxycarboxylic acid receptor 2 (HCAR2/GPR109A) may play an important role. HCAR2/GPR109A has been shown to regulate immune cell responses that potentiate anti-inflammatory signaling upon its activation in various tissues as evidenced principally by suppressed pro-inflammatory cytokine secretion in various experimental model systems. We have demonstrated HCAR2/GPR109A expression in RPE, microglia and endothelial cells and, our *in vitro* studies support that the receptor elicits anti-inflammatory signaling in these cell types. However, the functional relevance of HCAR2/GPR109A expression and its activation in the retina of the living animal has not been demonstrated definitively. This is the purpose of the present study.

**Methods:** Retinal function was evaluated temporally in wildtype (*Hcar2/Gpr109a ^+/+^*, WT) and knockout (*Hcar2/Gpr109 ^-/-^*, KO) mice via electroretinography (ERG). Fundoscopic imaging, spectral domain-optical coherence tomography (OCT), fluorescein angiography and post-mortem histological analyses were additionally performed to evaluate retinal health. Gene microarray, RT-qPCR studies, ingenuity analyses and proteome pathway mapping were performed to evaluate potential key differences in the molecular signatures of WT and KO mouse retinas. Leukostasis and flow cytometric assays were performed to demonstrate the *in vivo* impact of HCAR2/GPR109A expression and its therapeutic activation on pro-inflammatory immune cell trafficking in retina.

**Results:** Longitudinal studies revealed progressive anomalies in retinal morphology and function in HCAR2/GPR109A knockout mice that impacted the entire retina. Gene expression and protein interactome analyses revealed differences in gene and protein expression consistent with the increased immune reactivity and infiltration of bone-marrow derived immune cells detected in KO mouse retinas. Studies conducted in an acute model of retinal (endotoxin-induced) inflammation revealed that targeting the receptor via intraperitoneal administration of agonist, beta-hydroxybutyrate, limits immune cell activation, infiltration and related inflammation in WT retinas.

**Conclusions:** The present studies demonstrate a central role for HCAR2/GPR109A in regulating the complex interplay between resident retinal cells and peripheral immune cells and, the potential therapeutic utility that targeting the receptor holds with respect to preventing and treating inflammatory retinal diseases.

**Highlights:** Oxidative stress and inflammation are major causative factors in degenerative retinal diseases stemming from numerous causes (e.g., aging, diabetes, sickle cell). Thus, identifying new targets and developing strategies to counter these factors to prevent and treat retinal degeneration is important. The present in vivo study demonstrates convincingly the principal role of the hydroxycarboxylic acid receptor 2 (HCAR2/GPR109A) as a major regulator of retinal immune responses under normal conditions and therefore, as a target with extremely high potential for therapeutic modulation of these responses in retinal disease.

## INTRODUCTION

Inflammation is essential to tissue homeostasis, enabling tissues to limit the detrimental effects of infection and other cellular perturbations and persist functionally intact post-insult. When unregulated however, things often go dangerously awry. Indeed, chronic unregulated immune reactivity and consequent oxidative stress and inflammation is linked causally to the development and progression of a number of degenerative diseases. Retina is highly susceptible to damage as evidenced by the fact that low-grade but persistent inflammation is implicated heavily in the pathogenesis and progression of age-related macular degeneration (AMD) and diabetic retinopathy, leading causes of blindness worldwide. Thus, understanding better the mechanism(s) by which retina, a tissue traditionally characterized as “immune privileged” or otherwise isolated from systemic immunological perturbations, preserves the integrity of its physical barriers and coordinates endogenous and peripheral immune responses to protect itself from chronic inflammatory insult under normal physiologic conditions and, how to impact these mechanism(s) therapeutically in diseased conditions is of paramount importance.

The hydroxycarboxylic acid receptor 2 (HCAR2), also known as GPR109A was identified more than a decade ago as the G_i_-linked protein coupled receptor responsible for mediating the pharmacologic actions of nicotinic acid (14,21,26). Several years later, the ketone metabolite beta-hydroxybutyrate (BHB) and short-chain fatty acids like butyrate were identified as endogenous ligands (22–24). Outside of retina, HCAR2/GPR109A expression has been demonstrated in adipocytes, the cell type where the anti-lipolytic actions are most warranted, immune cells and in the epithelia of gastrointestinal tissues where it has been reported to play roles ranging from nutrient sensing to immune regulation (4,19,20,24). Regarding the latter function, HCAR2/GPR109A has been shown to potentiate responses that are collectively anti-inflammatory and/or anti-tumorigenic upon its activation by specific agonists like nicotinic acid, BHB and monomethylfumarate (5,16,20,23,24). In retina, we have demonstrated HCAR2/GPR109A to be expressed robustly in the basolateral membrane of the retinal pigment epithelium (RPE), microglia and endothelial cells (6,11,12). The former two cell types are principal components of the outer- and inner-blood retinal barriers, respectively, and therefore participate directly in the isolation of the retina from systemic insults. Furthermore, all three cell types secrete pro- and anti-inflammatory molecules in response to diverse stimuli and thereby participate directly in immunity at the molecular level, influencing the activation of local immune cells in retina and/or, the infiltration of immune cells located peripherally. Thus, with respect to the normal healthy retina’s known exceptional ability to regulate inflammation, these three cell types are key players. It is not surprising therefore, that these are the cell types in which HCAR2/GPR109A is expressed (6,8,13). So, while the precise mechanisms that govern barrier maintenance and immune responses in retina are not fully understood, it is highly plausible that HCAR2/GPR109A is involved.

Others and we have convincingly demonstrated a pattern of upregulated anti-inflammatory signaling and cellular protection in association with HCAR2/GPR109A activation in the retina (5-7,10,16). However, the functional significance of HCAR2/GPR109A expression in the intact retina normally or in pathologic conditions is still not whether understood. Given the devastating effects of chronic, unregulated inflammation in retina and the related potential impact that therapies aimed at enhancing the effects of potential existing anti-inflammatory signaling such as that which might be mediated via HCAR2/GPR109A might have with respect to prevention dysfunction and/or degeneration in this tissue, we sought to shed light on this topic by examining comprehensively the histologic, molecular and functional phenotype of wild-type and global HCAR2/GPR109A knockout mice maintained in normal housing conditions for up to 14 months. Further, we evaluated the impact of HCAR2/GPR109A expression and therapeutic augmentation of its activity in wild-type and HCAR2/GPR109A knockout mice exposed to acute pro-inflammatory stimuli. The current study demonstrates a central role for HCAR2/GPR109A both in preserving the highly selective retinal barriers and regulating the complex interplay between resident retinal cells and peripheral immune cells. Collectively, our findings support the potential utility of targeting the receptor therapeutically to prevent or treat inflammatory retinal diseases.

## Materials and Methods

### Animals

*Hcar2/Gpr109a*^−/−^ mice have been described previously (4). Heterozygous (*Hcar2/Gpr109a^+/−^*) breeding pairs of mice were used such that wildtype (*Hcar2/GPR109a^+/+^*) and knockout (*Hcar2/Gpr109a^+/−^*) mice were obtained from the same litter. The genotypes of the animals used in the study were confirmed by PCR using specific primer pairs. Normal male C57BL/6J mice were purchased from the Jackson Laboratory (Bar Harbor, ME) and used for age-matched comparative analyses. Mice were grouped by age as follows: young (1-3 months), middle (4-7 months) and advanced age (8-12 months). The care and use of all animals adhered to institutional guidelines for the humane treatment of animals and to the Association for Research in Vision and Ophthalmology (ARVO) for the Use of Animals in Ophthalmic and Vision Research.

### Quantitative Reverse Transcription Polymerase Chain Reaction (qPCR)

qPCR was used to quantify the steady-state levels of genes including C-C Motif Chemokine Ligand 2 (Ccl2), interleukin-1beta (IL-1β), tumor necrosis factor-alpha (TNF-α), and intercellular adhesion molecule 1 (ICAM-1) in mRNA samples isolated from wildtype and HCAR2/GPR109A knockout mouse retina/RPE obtained from control (untreated) mice or mice treated with PBS or LPS in the presence the HCAR2/GPR109A agonist beta-hydroxybutyrate (BHB); (n=6-9 male mice per group). Primer sequences available upon request. Hypoxanthine phosphoribosyltransferase 1 (HPRT1), 18S ribosomal RNA or glyceraldehyde phosphate dehydrogenase (GAPDH) were used internal controls for PCR reactions. Samples were run in duplicate and each RT-qPCR experiment was repeated at least three times. Real-time qPCR amplifications, using detection chemistry (SYBR Green; Applied Biosystems), were run in triplicate on 96-well reaction plates.

### Microarray analysis

Total RNA was prepared the retina (neuroretina and RPE combined) of male, wildtype (C57BL/6J) and *Hcar2/Gpr109a-*null mice eyes at 6 and 12 months (n=6 per genotype and age). Gene microarray analyses and associated Ingenuity Pathway Analyses (IPA) were performed following our established protocol (5).

### Electroretinography (ERG)

Visual function was assessed in young (2-5 months of age), middle-aged (6-9 months of age) and aged (10+ months of age) animals using electroretinography as per our published protocol (15). In brief, mice were dark adapted overnight, and then anesthetized using isoflurane anesthesia. Proparacaine HCl (0.5%, Akorn, Lake Forest IL), Tropicamide (0.5%, Akorn), and Phenylephrine HCl (2.5%, Paragon, Portland OR) eyedrops were applied for topical anesthesia and mydriasis. Stimuli were generated by a white LED. Light from the LED was presented to the eyes by 1 mm diameter optic fibers that were placed just in front of the pupils. Signals were acquired by silver-impregnated threads placed gently on the corneas, with a drop of hypromellose to keep eyes moist and enhance electrical contact. These signals were transferred to a Psylab amplifier (Contact Precision Instruments, Boston MA), with a gain of 10000, filtered between 0.3 and 400 Hz, with a notch filter at 60 Hz. The amplified signals were digitized by a National Instruments (Austin TX) 6323 data acquisition module, and read into custom software written in Igor Pro (WaveMetrics Inc, Lake Oswego OR). Stimulus intensity was calibrated in scotopic lumens. A series of increasing intensities of 5 ms flashes was presented while dark adapted. Amplitude and timing were measured for a-, b-, and c-waves, and a frequency domain analysis of the kernels derived by correlating the flash stimuli with the responses provided more global measures of amplitude and timing (18).

### Fundoscopy and Fluorescein Angiography

Fundoscopy and fluorescein angiography (FA) was performed to investigate the morphology of the retinal vasculature using Micron IV instrumentation (Phoenix Research Laboratories, Pleasanton, CA). Briefly, mice were anesthetized using ketamine/xylazine rodent anesthesia cocktail followed by dilation of the pupils with 1% tropicamide (Bausch and Lomb, Tampa, FL). Systane lubricant eye drops (Alcon, Ft. Worth, TX) were applied to maintain corneal moisture during the procedure. The mouse was positioned on the imaging stage for fundoscopy. For FA, the mouse was injected i.p. with 10–20 μl fluorescein sodium (10% Lite; Apollo Ophthalmic, Newport Beach, CA). Fluorescent images were then rapidly acquired at 30 s intervals for ∼5 min.

### Spectral Domain-Optical Coherence Tomography (SD-OCT)

SD-OCT studies were additionally performed on live wildtype and HCAR2/GPR109A knockout mice to monitor the integrity of the retina in cross-section also per our published method (15).

### Trypsin Digestion

Freshly enucleated eyes were fixed in 4% (wt/vol) paraformaldehyde in PBS overnight. After fixation, the retinas were dissected, washed in PBS and incubated in trypsin (0.5% (DifcoTrypsin 250) prepared in 20mM Tris buffer pH 8) for about 45 minutes at 37 °C. The vessel structures were isolated from the retinal cells by gentle shaking and rinsing in distilled water for overnight. After the vascular specimens were mounted on a slide and dried for 3-4 days at room temperature, periodic acid-Schiff staining was performed. Slides were images using traditional brightfield microscopy and the number of acellular capillaries per mm^2^ of capillary area was determined by counting 10 randomly selected microscopic fields.

### Immunofluorescence

Glial fibrillary acidic protein (GFAP) expression was analyzed in retinal cryosections (10 μm thickness). In brief, cryosections of mouse eyes were fixed in 4% paraformaldehyde for 10 min, washed with PBS, and blocked with Power Block (1X dilution) for 10 min at room temperature. Sections were then incubated overnight at 4 °C with rabbit anti-GFAP (1:500; AbCam) primary antibody. Retinal cryosections incubated with normal goat serum in the absence of primary antibody served as negative controls. Following multiple PBS washes, sections were incubated with secondary antibody, goat anti-rabbit Alexa Fluor Alexa Fluor 488 (1:1000), for 45 min at room temperature. Sections were rinsed again with PBS and incubated with Hoechst nuclear stain (1:10,000) for 5 minutes. Following this, sections were quickly rinsed with distilled water and coverslips were mounted using Fluoromount mounting media (Sigma, St Louis, MO). Additional cryosections from each group were also subjected to aminophenyl fluorescein staining per the manufacturer’s instructions (APF detection kit, ThermoFisher) to detect reactive oxygen species or, stained with hematoxylin and eosin for routine morphological analyses. RPE flatmounts were additional prepared and immunolabeled with anti-ZO-1 antibody as per our published method (Promsote et al., 2016).

### Lipopolysaccharide (LPS)-induced uveitis

Acute inflammation was induced in wildtype and HCAR2/GPR109A knockout mice (male, age 6 weeks) using LPS (*Salmonella typhimurium*, 4 mg/kg) (2). Wildtype and HCAR2/GPR109A knockout mice were injected intraperitoneally with PBS or β-hydroxybutyrate (BHB, 300mg/kg) once daily for three consecutive days. On day 4, LPS (4 mg/kg) was administered intraperitoneally in conjunction with PBS or with BHB (300 mg/kg) and, 24 h later some animals were subjected to leukostasis assay (as described below) and, others were sacrificed and eyes enucleated and processed for FACS analyses also as described below.

### Leukostasis Assay

To monitor the presence of adherent leukocytes in the retinal microvasculature of control (untreated) wildtype and HCAR2/GPR109A knockout mice, or, wildtype and HCAR2/GPR109A knockout mice treated as specified above (LPS-induced uveitis), leukostasis assays was performed as per our established protocol (1).

### Fluorescence activated cell sorting (FACS)

Freshly isolated retinal homogenates were suspended in PBS with 1% fetal bovine serum and stained with CD45, APC-CD41 (BD Biosciences), FITC-CD71 (BD Biosciences), Gr-1, and/or CD11b (BD Biosciences). Samples were analyzed using BD LSR II SORP and Diva 7.0 software (GRU Cancer Center Flow Cytometry Facility).

### Statistical Analysis

All experiments were repeated three to five times with independent tissue preparations and samples were run in duplicate. Data are presented as mean ± standard error of the mean (SEM). Statistical significance was determined with the Student t-test and one-way ANOVA with Tukey–Kramer’s post-hoc tests for comparisons between two groups or multiple groups, respectively. Differences were considered statistically significant at p<0.05.

## Results

### Retinal function is severely compromised in association with absence of HCAR2/GPR109A expression

To evaluate the functional relevance of HCAR2/GPR109A expression in the retina of the living animal, electroretinogram (ERG) recordings were obtained from young (2-5 months of age), middle-aged (6-9 months of age), and aged (10-12 months of age) wildtype (WT) and HCAR2/GPR109A knockout (KO) mice and used as an index of retinal function. A summary of scotopic flash response ERG recordings obtained from mice at 2, 5 and 12 months of age is presented in Fig. 1 (Note: ERG data obtained from wildtype and knockout mouse retinas have been plotted independently however, the axes scales used to graph the data are identical to facilitate accuracy in cross-comparison). As anticipated based on our prior longitudinal observations and those of others, a clear age-dependent decline in a-, b- and c-wave amplitudes was detected in wildtype mouse retinas (Fig. 1A, 1C and 1E, respectively). However, significant reductions in a-, b- and c-wave amplitudes were observed in knockout mouse retinas at all ages. ERG recordings from young, middle-aged and aged knockout were virtually indistinguishable (Fig. 1B, 1D, and 1F, respectively). Notably, ERGs obtained from young HCAR2/GPR109A knockout mice (2 months of age) were comparable to those of wild-type animals of advanced age (12 months), emphasizing the severity of the early retinal dysfunction present in knockout mice. The significance of the differences in ERG records obtained from wildtype versus knockout mouse retinas is highlighted in Figure 1G, in which ERG data obtained from middle-aged wild-type (red) and HCAR2/GPR109A knockout mice (blue) has been isolated and presented together in the same graph. Collectively, ERG data suggest that pronounced functional alterations in retinal neuronal (photoreceptor and bipolar cells) and pigment epithelial cells, the cells responsible for generating the a-, b- and c-waves, respectively, occur early and persist throughout aging in association with loss of HCAR2/GPR109A expression. To determine whether specific cellular layers and/or populations of affected cells could be identified at the morphological level to support the functional abnormalities revealed by ERG, we next performed detailed *in vivo* imaging studies and post-mortem histologic analyses.

**Figure 1.**
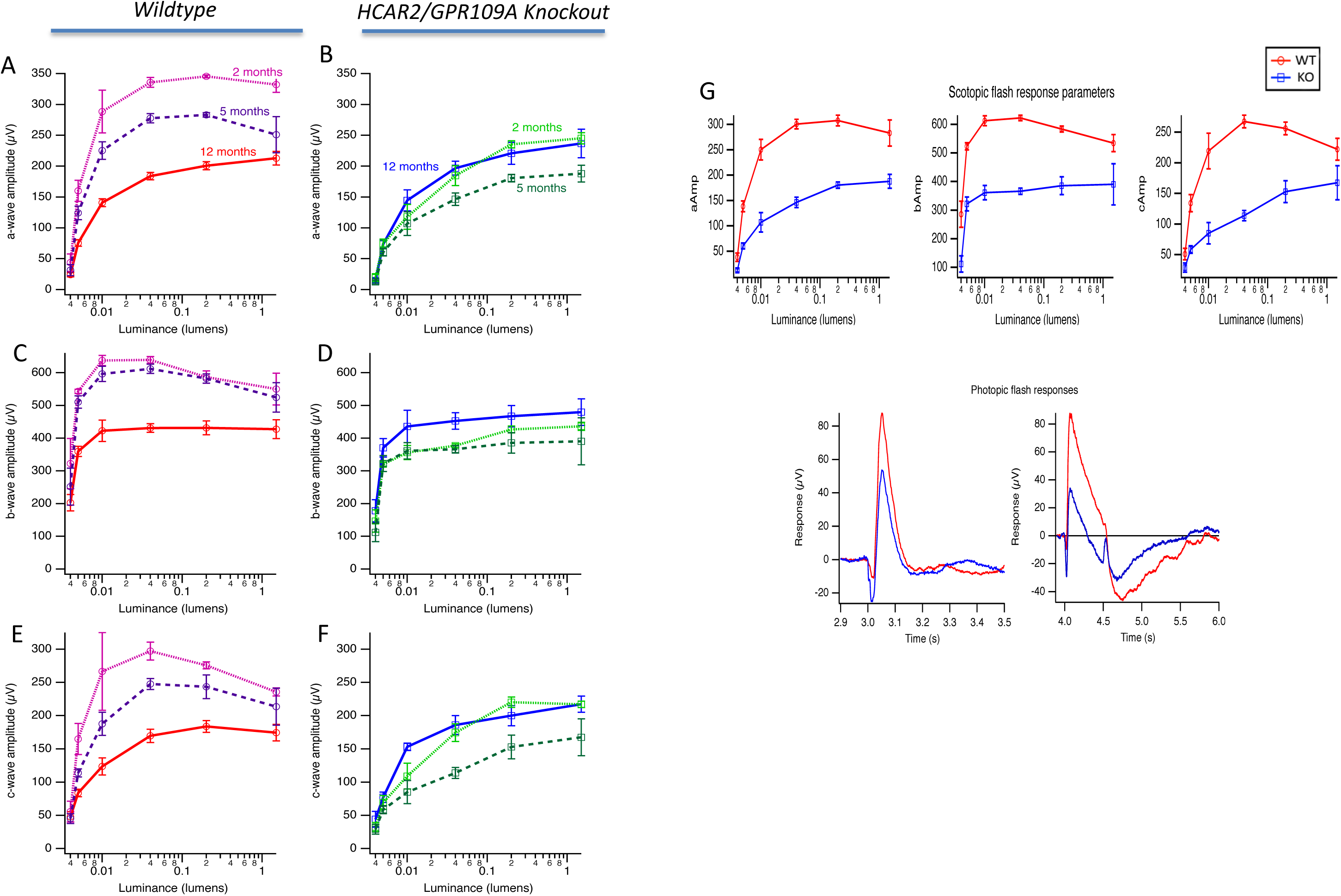
Early retinal dysfunction in HCAR2/GPR109A knockout mouse retinas as revealed by electroretinography (ERG). ERG studies were performed on male, wildtype and HCAR2/GPR109A knockout mice at 2, 5 and 12 months of age (n=6-9 mice per group). Age-dependent reductions in a-, b- and c-wave amplitudes (Figs. 1A, 1C, and 1E, respectively) were apparent in wildtype animals. However, these same ERG components (Figs. 1B, 1D, and 1F, respectively) were virtually indistinguishable in HCAR2/GPR109A knockout mice irrespective of age. ERG data from wildtype and HCAR2/GPR109A knockout mice at 5 months of age are compared directly in Fig. 1G to highlight the pronounced differences in retinal function that exist amongst these groups.

### Loss of HCAR2/GPR109A results in progressive retinal morphological changes and disruptions

Spectral domain-optical coherence tomographic (SD-OCT) imaging was performed on young (Fig. 2A-B), middle-aged (Fig. 2C-D) and aged (Fig. 2E-F) wildtype and HCAR2/GPR109A knockout mice. SD-OCT images obtained from wildtype eyes were largely unremarkable at all ages (Fig. 2A, 2C, 2E). The organization of the wildtype retina was normal and non-disrupted, the signal-to-noise ratio was appreciably high and therefore, the associated clarity of the retinal images obtained was good. Alternately, signs of overt pathology were detected in the eyes of most HCAR2/GPR109A knockout mice. Subtle changes were detected even at very young ages (Fig. 2B). Additionally, the clarity of SD-OCT images obtained from knockout mouse eyes was often reduced either due to apparent edema and/or the presence of obstructions, presumably cellular debris; both of which are apparent in Fig. 2F (upper left panel). In a number of instances, the abnormalities present in the vitreous were sizable enough to impede the proper passage of light as evidenced by darkened regions of retina which indicate regions of tissue that could not be visualized with reasonable clarity. The same was detected in the eyes of some younger knockout mice, although less commonly (Fig. 2B, red arrow). With increasing age, abnormal vascular growth also became an increasingly common finding in knockout mouse eyes (e.g., cross-section of an artery in Fig. 2D, left panel, white arrow); vitreoretinal membrane formation could also be appreciated (Fig. 2F, lower left and right panels, white arrows). Collectively, SD-OCT findings in aged HCAR2/GPR109A knockout mouse eyes are consistent with edematous changes and potential robust inflammation. As expected in live animals, there was some variability in our imaging findings. Therefore, it is important to note that the images that we have presented in Figure 2 and beyond are representative of the most commonly observed wildtype and knockout mouse retinal phenotypes, not the most unique or severe. This variability is corroborated by associated morphometric data obtained during SD-OCT assessments (Figs. 2G-O) in which few statistically significant differences were identified.

**Figure 2.**
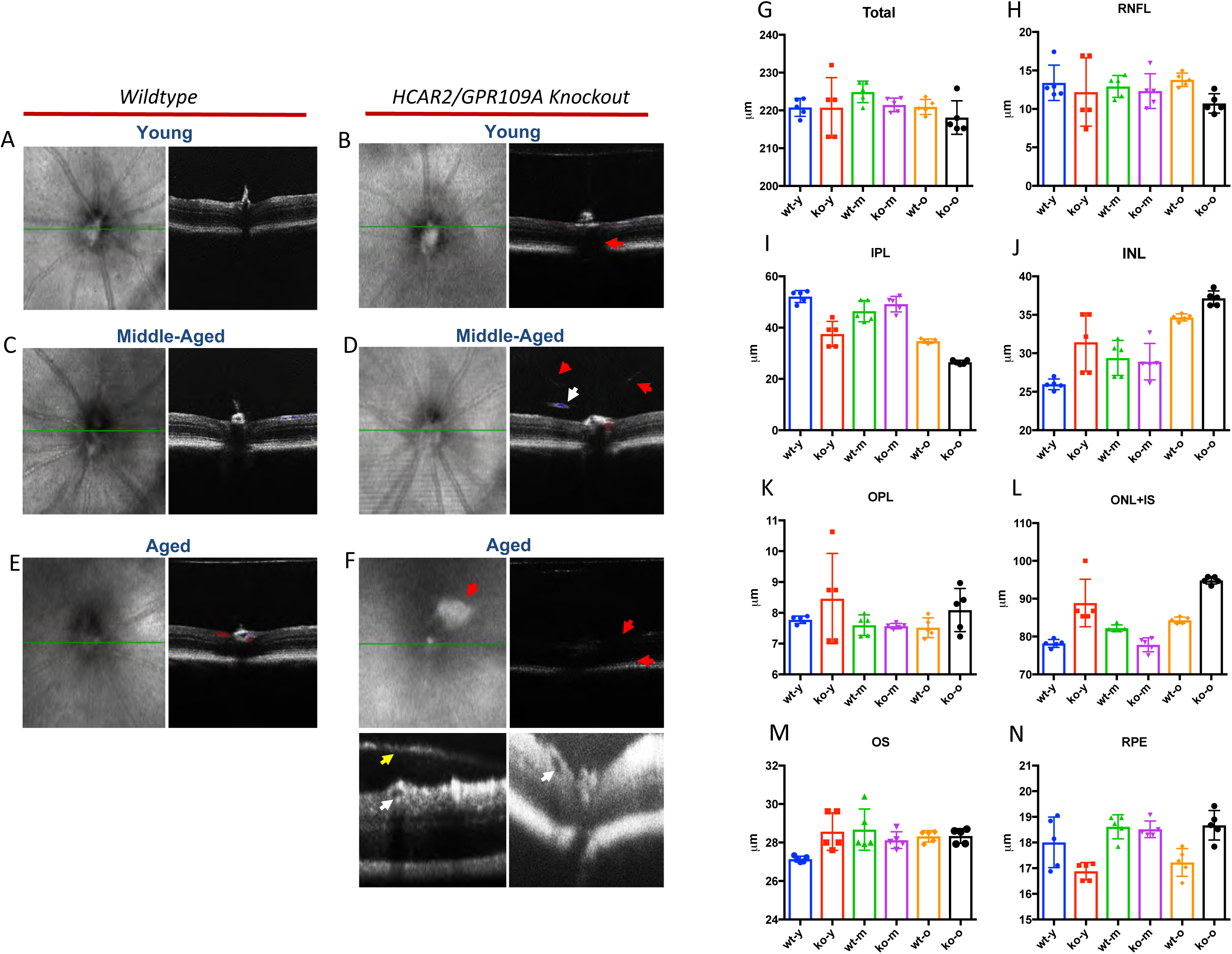
Spectral domain-optical coherence tomography (SD-OCT) imaging of wildtype and HCAR2/GPR109A knockout mouse retina. SD-OCT imaging of the central retina was performed on anesthetized young (2-4 months of age), middle-aged (5-9 months of age) and aged (10-14 months of age) wildtype (Figs. 2A, 2C, and 2E) and HCAR2/GPR109A knockout mice (Figs. 2B, 2D, and 2F). n=6-9 male mice per group. SD-OCT-guided morphometric analyses to include measurements of total retinal thickness (Fig. 2G), nerve fiber layer thickness (Fig. 2H), inner plexiform layer thickness (Fig. 2I), inner nuclear layer thickness (Fig. 2J), outer plexiform layer thickness (Fig. 2K), outer nuclear layer thickness (Fig. 2L), outer segment thickness (Fig. 2M) and retinal pigment epithelial (RPE) thickness (Fig. 2O) were additionally obtained. Data are presented as mean +/- standard error and *p*<0.05 was considered statistically significant.

### Compromised retinal and vascular integrity in HCAR2/GPR109A knockout mouse retina

Though useful in gauging phenotypic differences in HCAR2/GPR109A knockout mouse eyes compared to wildtype, SD-OCT imaging studies provided little information relevant to the integrity of the vascular components of the retina in HCAR/GPR109A mice. Thus, to determine whether morphological abnormalities associated with the absence of HCAR2/GPR109A expression extend beyond the neuroretina and RPE, fundoscopic and fluorescein angiographic (FA) imaging of live mouse eyes was performed (Fig. 3A). Retinal fundus photographs obtained from wildtype mice were largely unremarkable at all ages observed as indicated by the representative fundus image obtained from an aged wild-type mouse shown in the upper left panel of Fig. 3A. The vascular phenotype of wild-type animals was also generally intact and uncompromised as per FA images (Fig. 3A, upper right panel). There was no evidence of extravascular fluorescein dye or vessels that were overly dilated and/or tortuous in appearance. Fundus and FA images of young HCAR2/GPR109A knockout mouse retinas were highly similar to those of wildtype at comparable age and therefore also generally unremarkable (data not shown). However, slightly translucent areas (demarcated by hashed yellow lines in Fig. 3A), indicative of possible retinal thinning and/or hypo-pigmentary changes in the RPE, were evident in many fundus images obtained from HCAR2/GPR109A knockout mice of middle and advanced age (Fig. 3A, left middle and left bottom panels, respectively). FA imaging of the retinas of these same knockout mice revealed the presence of abnormally dilated and/or unusually tortuous central retinal vessels (Fig. 3A, red arrows). Interestingly, arteriovenous communications and vascular nicking or collaterals, extremely rare findings in normal mouse retina, were detected in some knockout mouse eyes (Fig. 3A, middle right panel, white asterisks and yellow arrows, respectively), as were large areas of extravascular fluorescein synonymous with vascular leakage that appeared to correlate closely with some of the translucent areas that were evident in fundus images (Fig. 3A, lower right panel, white hashed circle). Fluorescein leakage was also prominent at the optic disc in animals of advanced age (Fig. 3A, lower right panel, red hashed circle). The disrupted vascular phenotype detected in FA imaging of HCAR2/GPR109A knockout mouse retinas is corroborated by trypsin digest preparations of wildtype and knockout mouse retinas which revealed the disorganized arrangement of vessels in HCAR2/GPR109A knockout mouse eyes as well as a high number of degenerate capillaries (Fig. 3B, black arrows), findings that were not common to wildtype retinas of comparable age. Collectively, the vascular anomalies detected in HCAR2/GPR109A knockout mouse eyes are congruent with the existence of a retinal environment in these animals that is inflammatory and potentially hypoxic; a finding supported by hydroxyprobe (anti-pimonidazole hydrochloride) and anti-isolectin b4 co-immunolabeling of retinal flatmounts prepared from young, wildtype and knockout mouse eyes (Fig. 3C). Areas of tissue hypoxia are denoted by green fluorescence relative to the retinal vasculature (red fluorescence). Hypoxic areas were detected throughout HCAR2/GPR109A knockout mouse eyes and these areas increased in number and size proportionate to increasing age. Positive hydroxyprobe labeling was particularly intense in the region of the optic disc as shown in Fig. 3C (right panel). Alternately, minimal positivity (green fluorescence) was detected in retinal flatmounts prepared from wild-type mouse eyes.

**Figure 3.**
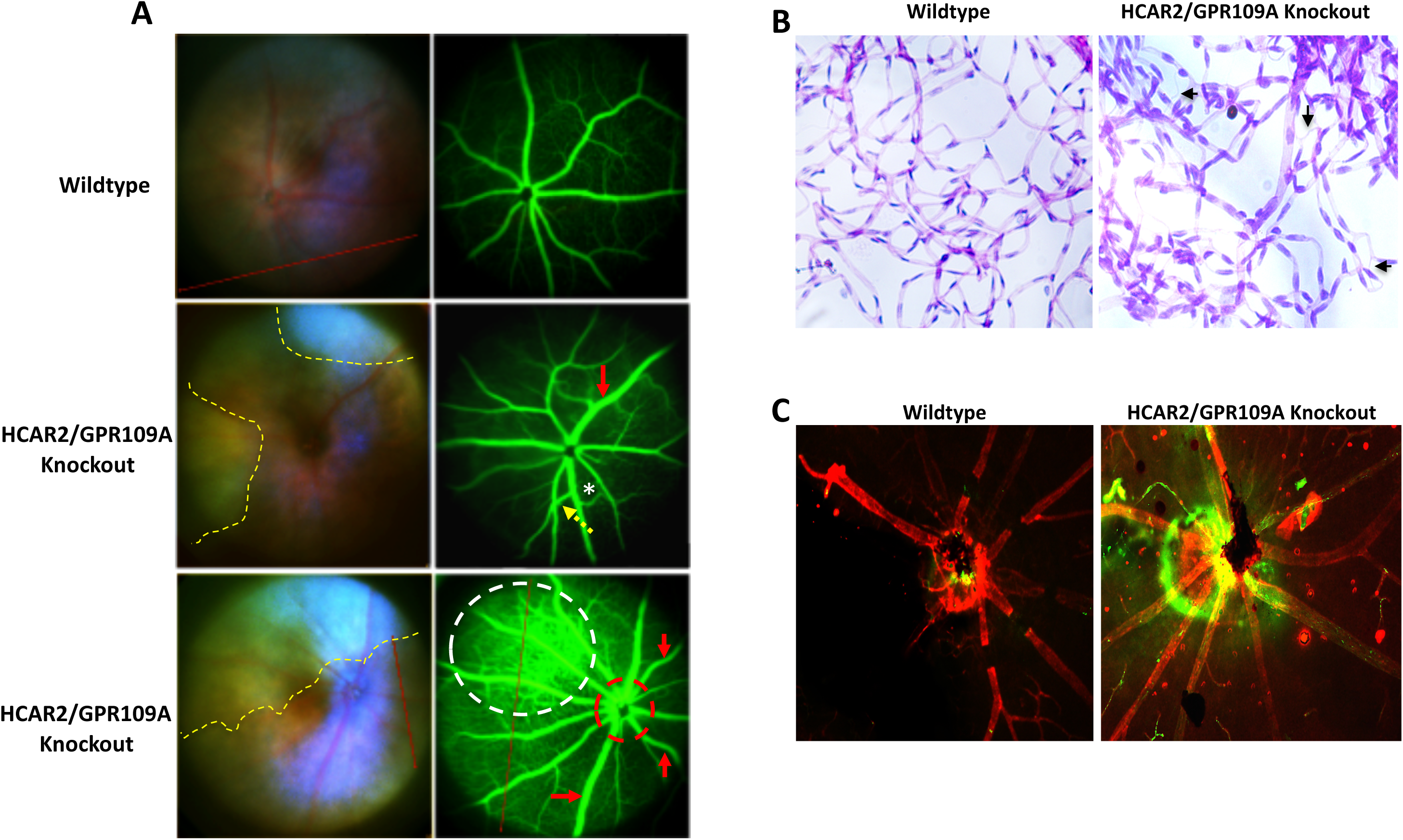
Fundoscopy and fluorescein angiography imaging of wildtype and HCAR2/GPR109A knockout mouse retina. Fundoscopic and fluorescein angiographic images were obtained from young (2-4 months of age), middle-aged (5-9 months of age) and aged (10-14 months of age) wildtype and HCAR2/GPR109A knockout mice Representative fundoscopy and fluorescein angiography images from aged wildtype (Fig. 3A, *top left and right panels, respectively*), middle-aged HCAR2/GPR109A knockout (Fig. 3A, *middle left and right panels, respectively*) and aged HCAR2/GPR109A knockout (Fig. 3A, *lower left and right panels, respectively*) mouse eyes are provided. Fundus and fluorescein angiography images of wildtype retinas were largely unremarkable at all ages observed with respect to the presence of overt pathology. However, evidence of potential retinal thinning (areas denoted in fundus images by yellow hashed lines) were evident in many HCAR2/GPR109A knockout fundus images. Additionally, progressive vessel abnormalities such as excessive dilation and tortuosity (*red arrows)*, vessel nicking or cross-over (*yellow arrow*) and/or potential arteriovenous anastomosis (*white asterisk*), leaking at the optic disc (*red hashed circle)* and peripheral retinal leakage (*white hashed circle)* were apparent in HCAR2/GPR109A knockout mouse eyes. Trypsin digest preparations were prepared from wildtype and HCAR2/GPR109A knockout mouse retinas. A high number of degenerate capillaries (*black arrows)* were noted in HCAR2/GPR109A knockout mouse retinas as indicated in the representative image provided in (B). (C) Representative images of hydroxyprobe-labeled retinal flatmounts reveal minimal positivity (*green fluorescence*) in wildtype retinas but particularly robust signal in HCAR2/GPR109A knockout mouse retinas especially close to the optic disc.

### Increased inflammation and oxidative stress in HCAR2/GPR109A knockout mouse retina

Our prior cell and molecular studies support a key role for HCAR2/GPR109A in the regulation of inflammatory signaling in retina (6) and, we now know that loss of HCAR2/GPR109A expression has a significant detrimental effect on retinal morphology and function. To confirm whether the functional and morpho-histological changes seen in HCAR2/GPR109A knock out mouse retinas are consistent with inflammation and increased oxidative stress as previously speculated, glial fibrillary acidic protein immunoreactivity and aminophenyl fluorescein (APF) staining, respectively, were performed on wildtype and HCAR2/GPRGPR109A knockout mouse cryosections (Fig. 4). Minimal GFAP expression was detected in wildtype mouse retinas (Figs. 4A-C, left panels). However, GFAP immunoreactivity was prominent in HCAR2/GPR109A knockout mouse retinas, particularly in those retinas obtained from young and middle-aged knockout mice (Figs. 4A-C, right panels). GFAP immunostaining was still evident but reduced in the retinas of HCAR2/GPR109A knockout mice of advanced age (Fig. 4C, right panel) compared to young and middle-aged HCAR2/GPR109A knockout mice. Importantly however, the retinas of these aged knockout animals were obviously disrupted as evidenced also by the reduced amount nuclear counter stain (blue immunofluorescence) present. Hence, the reduction in GFAP labeling in this group may be due more to the increased amount of tissue damage and consequent cellular loss than to an actual reduction in inflammation/GFAP immunoreactivity. Congruent with a robust inflammatory environment, aminophenyl fluorescence (APF) labeling revealed that levels of oxidative stress were too increased in HCAR2/GPR109A knockout mouse retinas. As reported in other studies of normal aging, oxidative stress (APF labeling, green fluorescence) increased relative to age in wildtype mouse retinas (Figs. 4D-F, left panels). However, a considerably higher amount of APF labeling was detected in HCAR2/GPR109A knockout mouse retinas compared to wildtype, especially in the young and middle-aged groups (Figs. 4D-E). Interestingly, APF staining localized most robustly to the outer retina of HCAR2/GPR109A knockout mice. Further, as noted with GFAP staining (above), less APF staining was evident in aged HCAR2/GPR109A knockout mouse retinas (Fig. 4F) again, likely attributable to the fact that a considerable amount of tissue damage and cellular loss had already occurred by this time.

**Figure 4.**
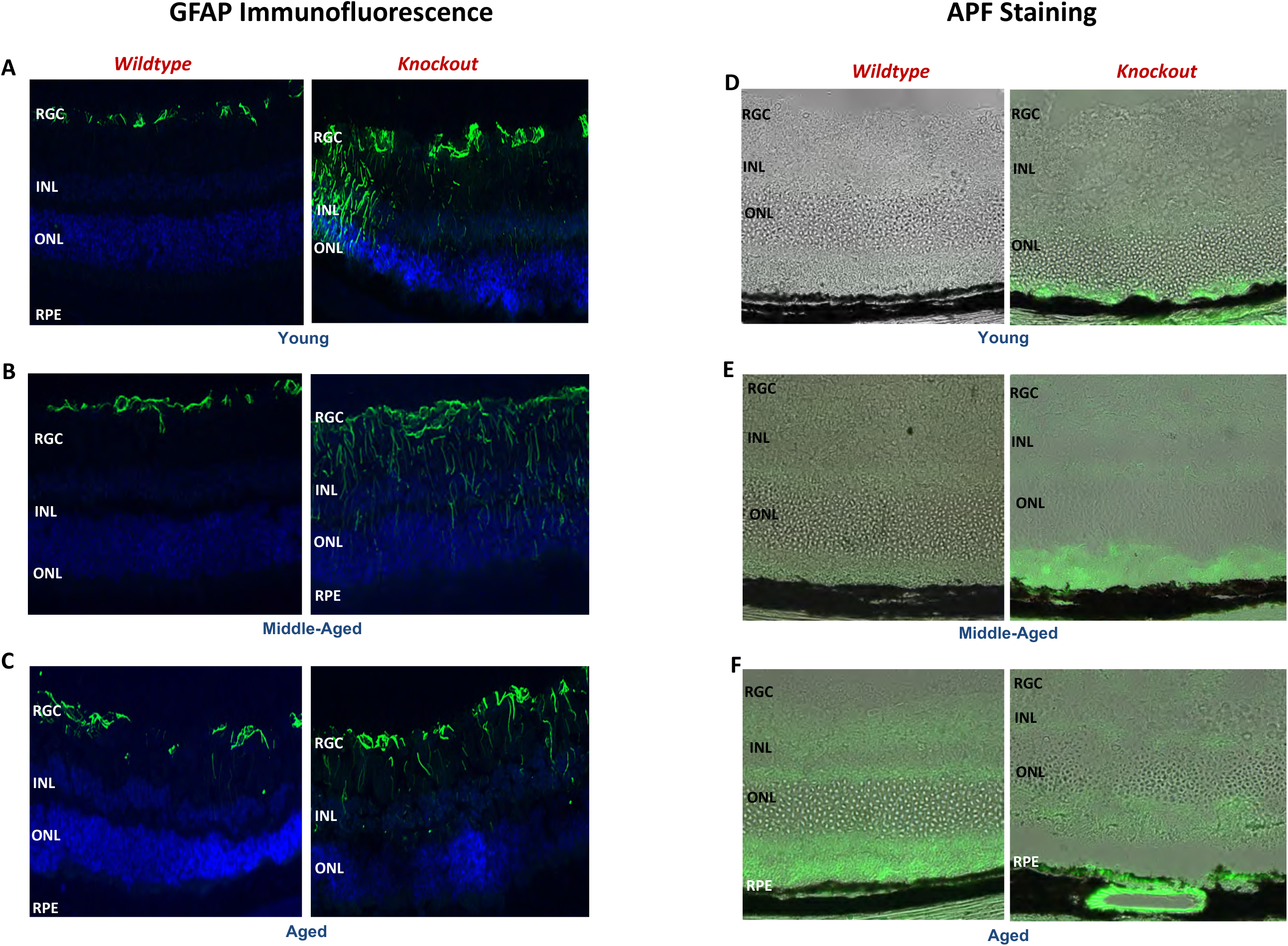
Evaluation of glial cell immunoreactivity and reactive oxygen species (ROS) in wildtype and HCAR2/GPR109A knockout mouse retina. (*A-C*) Glial cell immunoreactivity was monitored in retinal cryosections prepared from young (2-5 months, *A)*, middle-aged (6-9 months, *B*) and aged (10+ months of age) wildtype and HCAR2/GPR109A knockout mice via glial fibrillary acidic protein (GFAP) immunolabeling. (*D-F*) ROS production was also monitored in additional cryosections obtained from young (2-5 months, *D)*, middle-aged (6-9 months, *E*) and aged (10+ months of age, *F*) via aminophenyl fluorescein (APF) staining.

### Loss of HCAR2/GPR109A expression results in progressive retinal degeneration

Data obtained to this point support potential widespread functional and morphological disruption of retina congruent with the absence of HCAR2/GPR109A expression. To follow-up on this definitively and determine whether specific cellular layers and/or populations of affected cells could be identified at the histologic level to support the functional and morphological abnormalities identified, hematoxylin and eosin-stained retinal cryosections were prepared from wildtype and knockout animals of young, middle and advanced age and evaluated via traditional light microscopy for evidence of gross morphological alteration (Fig. 5). The well-organized structure of the neuroretina and RPE could be appreciated in wild-type mice of all ages. Alternately, a progressive degenerative phenotype was observed in HCAR2/GPR109A knockout mouse retinas. The organization or the retinal layers was increasingly disrupted with age and loss of cellularity was prominently detected in the ganglion cell layer (GCL), inner nuclear layer (INL) and outer nuclear layer (ONL) of most HCAR2/GPR109A knockout mice including those of relatively young age (Figs. 5B, 5D, 5F; *black arrows*). Pigmentary abnormalities and breaks in the basement membrane were additionally notable and detected frequently (Figs. 5B, 5D, 5F; *white arrows*). Further, choroidal thickening was prominently evident in HCAR2/GPR109A knockout mouse retinas from middle-age onwards (Figs. 5D, 5F; *brackets)*. The alterations in HCAR2/GPR109A knockout mice penetrated the entire retina and were not restricted to any particular region. To better highlight this and the notable differences in the organization of the inner (yellow brackets) and outer segments (white brackets) and the RPE/choroid (white arrows and circle) of HCAR2/GPR109A knockout mice, additional cryosections were prepared (Fig. 5G-I). These additional sections were stained using an adjusted hematoxylin and eosin-staining procedure in which the processing time for eosin staining was extended from 30-45 seconds to 1.0-1.5 minutes to allow deeper penetration of the dye and better resolution of the non-nuclear areas of the tissues.

**Figure 5.**
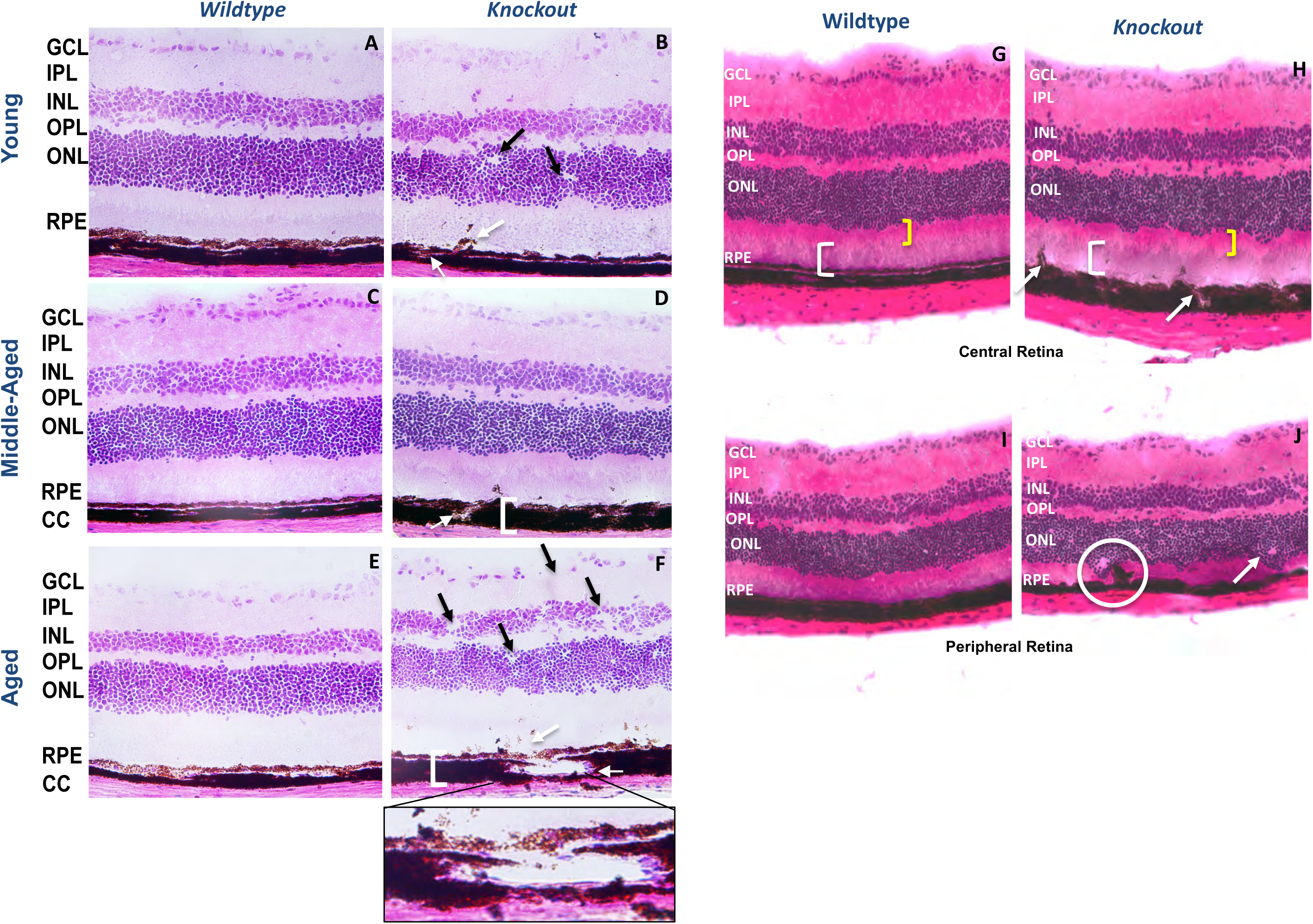
Morphological characterization of wildtype and HCAR2/GPR109A knockout mouse retina. Hematoxylin and eosin-stained (H&E) cryosections were prepared from wildtype and HCAR2/GPR109A knockout mouse retinas of various age. Representative images of young (*A* and *B*), middle-aged (*C* and *D*) and aged (*E* and *F*) wildtype and knockout images are provided. White arrows denote abnormal RPE-choroid communications. Black arrows point to areas of obvious cellular dropout in the nuclear layers, and white brackets in *D* and *F* denote choroidal thickening. Further, the callout in panel *F* highlights an area of significant RPE hypopigmentation and/or cellular loss immediately above a break in Bruch’s membrane and an abnormally large vessel in the thickened choroidal space; features common to retinas of HCAR2/GPR109A knockout mice of advanced age. Additional images of central and peripheral wildtype and knockout retina are provided that were obtained following use of a modified H&E-staining method (*G-J*). White brackets in panels *G* and *H* highlight the outer segments and inner segments, respectively such that differences in the thickness and overall morphology of these regions can be better appreciated. The white arrow in panel *H* points to a basement membrane break and abnormal RPE-choroid interface whereas, the white arrow in panel *J* points to a region of disrupted morphology in the outer nuclear layer. A similar morphological disruption involving both the RPE and outer nuclear layer is noted by the white circle in this same retinal section.

### HCAR2/GPR109A regulates outer retinal barrier function

Prior work by others and we demonstrate the robust expression of HCAR2/GPR109A in RPE. Additionally, studies above show convincingly that outer retina and RPE are prominently affected, functionally and morphologically, in the absence of HCAR2/GPR109A expression. To evaluate the impact of HCAR2/GPR109A expression in RPE more closely, RPE flatmounts were prepared from wildtype and HCAR2/GPR109A knockout mouse eyes and immunofluorescence analyses of zonula occludens-1 (ZO-1) were performed to evaluate the integrity of the RPE. Representative RPE flat mount images from wildtype and HCAR2/GPR109A knockout mouse retinas (age 10 months) are provided in Figure 6A. Flatmount preparations of from wildtype mice showed the characteristic cobblestone pattern of ZO-1 labeling (green fluorescence) consistent with intact RPE junctions and a well-functioning barrier. In contrast, ZO-1 staining of HCAR2/GPR109A knockout RPE flatmounts revealed areas of intact ZO-1/cobblestone-shaped RPE borders adjacent to areas of obvious disruption and diffuse ZO-1 labeling. This was the case, in 80-90% of all the HCAR2/GPR109A flatmounts evaluated (n=9). Therefore, we were unable to perform typical quantitative measures (e.g., quantification of total RPE cell number or the number of multinucleate RPE cells per a defined surface area). Given the striking degree of ZO-1 disruption in knockout retinas, we evaluated ZO-1 expression by qPCR in younger animals. A significant decrease in ZO-1 mRNA expression was obvious in animals as early as 5 months of age (Fig. 6B). Collectively, these data support a progressive deterioration of RPE barrier properties with age. To determine whether the absence of HCAR2/GPR109A expression affects cell monolayer permeability at the functional level, primary RPE cells were prepared from wildtype and HCAR2/GPR109A knockout mouse eyes per our established method (11) and grown for 6 weeks to confluency on transwell inserts and subjected to an *in vitro* cell monolayer permeability assay (15, 17). As anticipated based on morphological observations, barrier integrity was significantly reduced, as indicated by increased FITC-Dextran leakage, in HCAR2/GPR109A knockout primary RPE cells compared to wildtype RPE (Fig. 6C).

**Figure 6.**
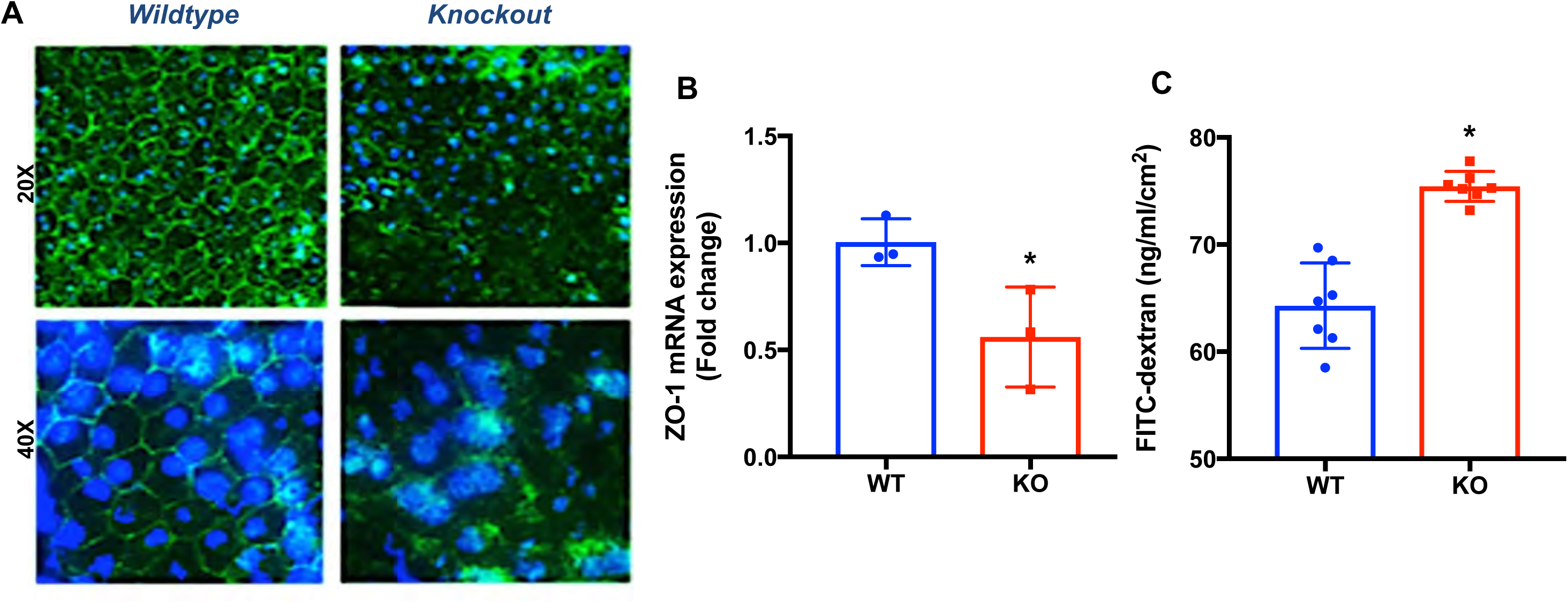
Analysis of outer blood-retinal barrier (retinal pigment epithelial, RPE) morphology and function in HCAR2/GPR109A knockout mouse retina. (*A*) RPE flatmounts were prepared from wildtype and HCAR2/GPR109A knockout mouse eyes and immunolabeled with anti-zonnula occludens 1 (ZO-1) to evaluate junctional complex formation. (*B*) Quantitative reserve transcriptase polymerase chain reaction (qRT-PCR) was performed to evaluate ZO-1 expression at the mRNA level. (*C*) *In vitro* permeability assays were performed using confluent monolayers of primary RPE cells isolated from wildtype and HCAR2/GPR109A knockout mouse eyes and grown on transwell supports. *p<0.01.

### HCAR2/GPR109a regulates immunomodulatory interactions in mouse retina

Our past studies support the importance of HCAR2/GPR109A in regulating inflammation and our present study demonstrates the impact of receptor loss on retinal morphology and visual function in the living animal however, the underlying mechanisms are still not fully understood. Thus, we sought to detect potential differences at the transcriptional level with respect to potential early and late gene changes that might explain the functional and morphological phenotypes that we have observed in HCAR2/GPR109A knockout mouse retina. Microarray analyses were performed using total RNA prepared from whole retina (neuroretina and RPE tissues). A heat map depicting gene expression changes detected in 6-month old male, wildtype versus knockout mice is provided in Figure 7A. Using a cut-off value of 1.5-fold expression minimum fold-change and a p-value <0.05, absence of HCAR2/GPR109A expression resulted in the differential expression of 136 transcripts. Of these, 29 transcripts exhibited 1.5 - 34-fold higher expression (red color) whereas 136 transcripts revealed 1.5 - 34-fold reduction (blue color). Using the same parameters to evaluate samples obtained from aged wildtype and HCAR2/GPR109A knockout mice, 96 genes were differentially expressed; a total of 82 genes were up-regulated (1.5 - 39.6-fold change) while 14 targets were down-regulated (1.5 - 52.4-fold change). This shows that the HCAR2/GPR109A knockout has distinct gene regulatory patterns and that these patterns differ relative to age. Though the fold-differences in expression varied, gene microarray data obtained from neuroretina/RPE samples of animals of different age was quite similar with respect to differentially expressed genes that were identified. Ingenuity® (Ingenuity Systems, Red Wood City, CA) pathway analyses of the gene microarray data revealed functional networks containing genes relevant to the top functions affected by knockout of the receptor in the retina: (1) cellular immune response, (2) reactive oxygen species generation and antioxidant signaling, and (3) the migration of bone-marrow derived (myeloid) cells (Figure 7B). A number of gene products were associated with each of these functional hubs, linking them in the network topology. Genes relevant to immune regulation accounted for the highest number of genes that were differentially expressed in HCAR2/GPR109A knockout mouse retina with 63.3% of the top transcripts having a known role in innate immunity or inflammation. Relatedly, the top canonical pathways identified involved immunomodulatory functions such as notch signaling (p=0.032) and amyloid processing (p=0.05) along with immune cell adhesion and movement (p=0.005). Other top pathways such as Notch, ErbB4(p=0.05), and amyloid processing are interconnected in that the processing of molecules within these pathways are carried out by γ-secretase cleavage. Calcium signaling (p-value=0.005) and superoxide radical formation (p-value=0.007) are both master signaling regulators that are relative to a number of the top pathways found in the analysis. Integrin-linked kinase (ILK) signaling (p=0.006) and HIPPO (p=0.07, p=0.007) were also identified as major pathways, and both involve cell proliferation and signal transduction. Top differentially regulated genes identified by microarray analysis were validated by qPCR (Table 1). A protein interactome map was additionally developed using established using protein-protein interaction map software and databases. This interactome map suggests that the changes in gene expression identified in the gene microarray translate to relevant differences in protein expression and interaction and therefore potential function with respect to relationship between signaling mediated by HCAR2/GPR109A and the regulation of retinal immune responses.

**Figure 7.**
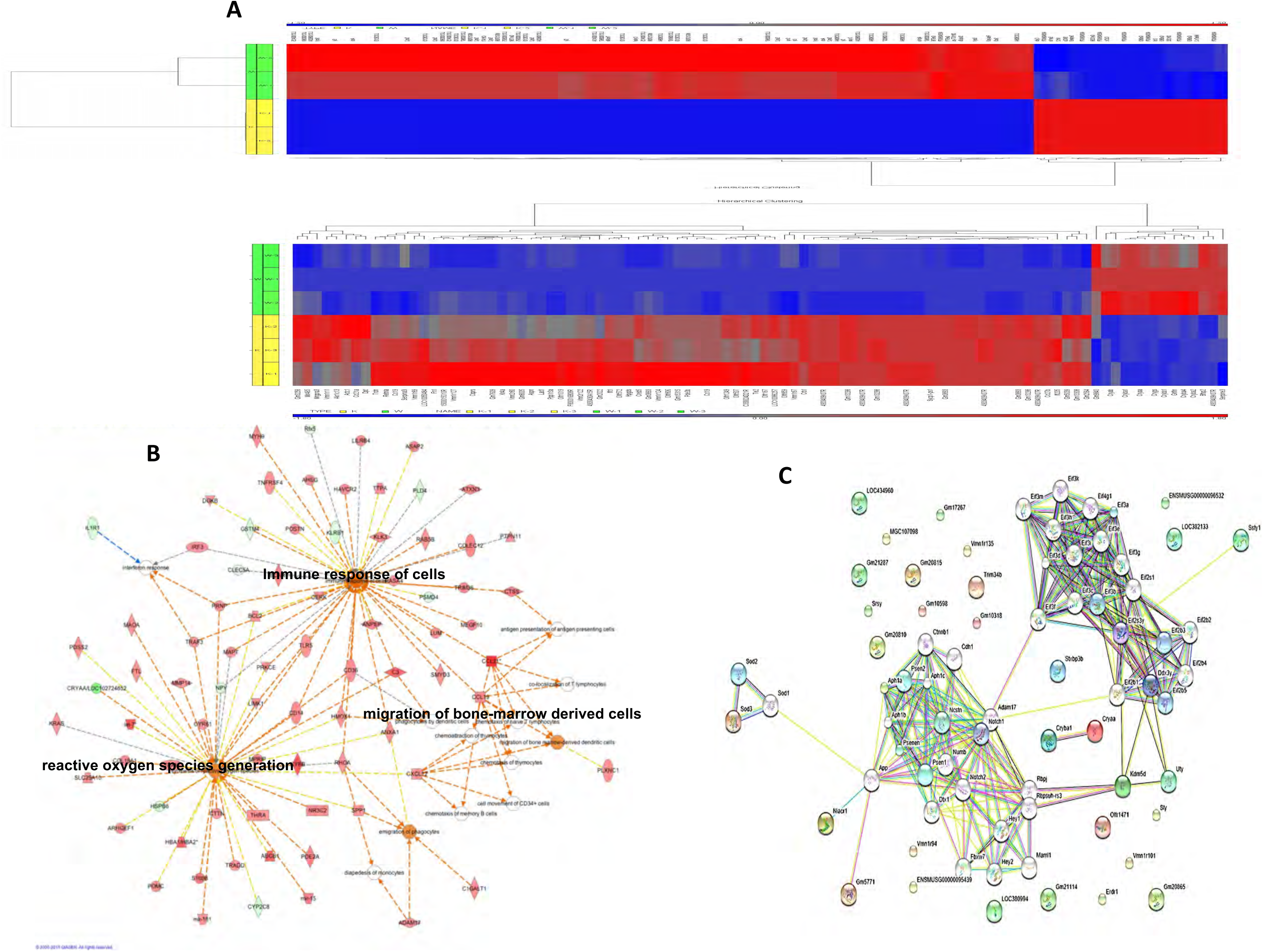
Microarray analysis of differentially regulated genes, gene networks and protein-protein interactions in wildtype and HCAR2/GPR109A knockout mouse neuroretina and RPE. RNA samples from wildtype and HCAR2/GPR109A knockout mouse neuroretina and RPE of various age were obtained and used for gene microarray analysis software. (*A*) A number of genes were differentially regulated in HCAR2/GPR109A knockout mouse eyes compared to wildtype as illustrated by the heat map derived from gene analyses conducted on male, 6-month old wildtype and HCAR2/GPR109A knockout mouse RNA samples. (B) Ingenuity pathway analyses were performed to explore the relationships between the genes most highly altered. (Green=genes downregulated in KO vs. WT, Red = genes upregulated in KO vs. WT). Genes belonging to functional or causal network pathways relevant to the immune response of cells, immune cell migration and reactive oxygen species generation were significantly impacted in HCAR2/GPR109A knockout mouse eyes and therefore ranked highest among shared pathways in which genes revealed to have the highest-fold expression with respect to differential gene expression between HCAR2/GPR109A knockout and wildtype mouse eyes. (C) Protein interactome mapping supported the findings of gene microarray and ingeniuty analyses with respect to the identification of functional networks relevant to differentially regulated genes.

**Table 1.**
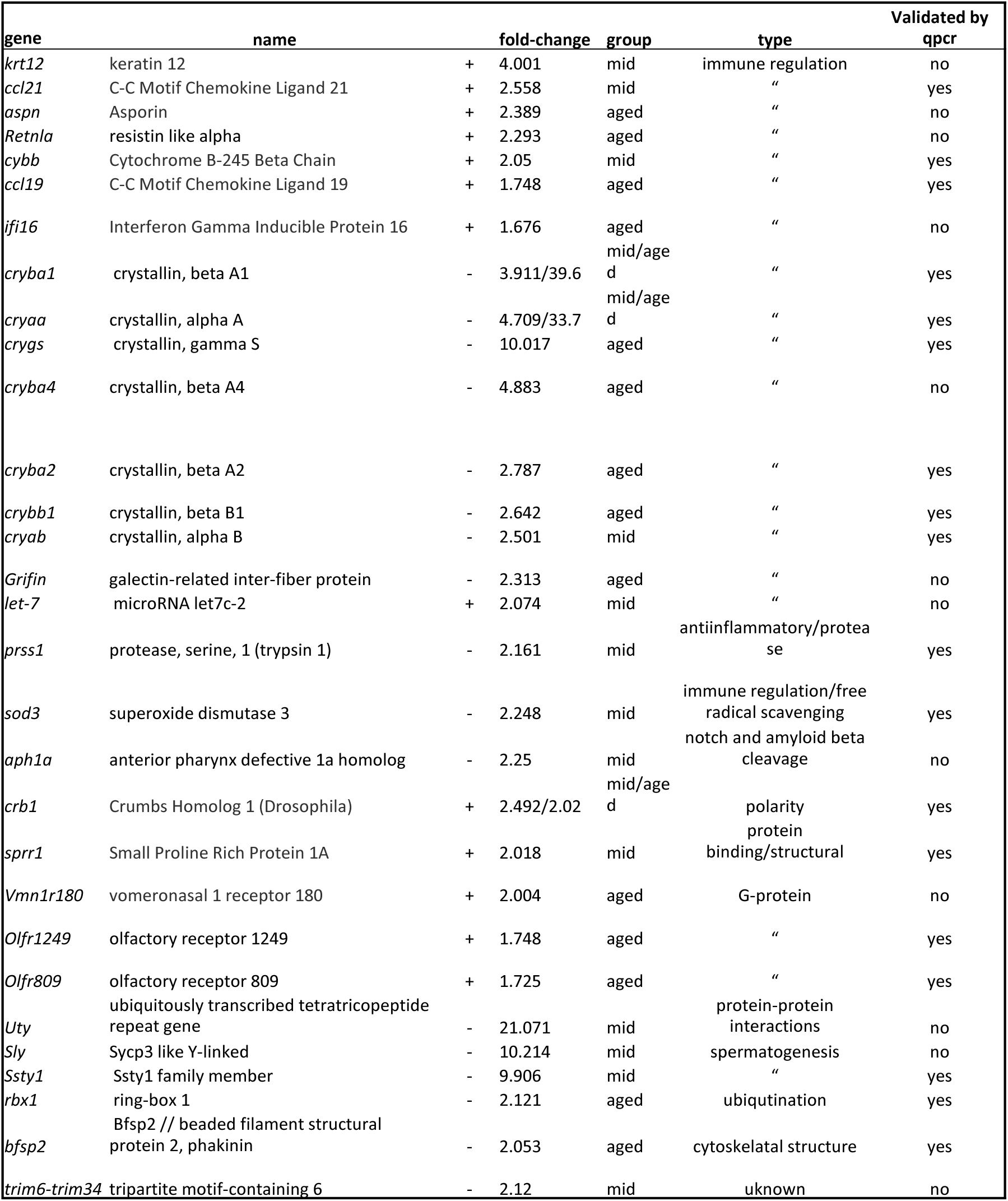
qRT-PCR validation of differentially regulated genes in wildtype (WT) and HCAR2/GPR109A knockout (KO) neuroretina/RPE eyecup.

### Confirmation of increased activation and infiltration of inflammatory immune cells in HCAR2/GPR109A mouse retina

Our prior published studies demonstrate a role for HCAR2/GPR109A in anti-inflammatory signaling in retina/RPE. Our present results provide definitive cues relevant to the molecular mechanisms that underlie this function, with regulation of immune cell migration and activation being paramount. Thus, we sought next to validate this in the living animal. To do so, we evaluated leukocyte adhesion in flatmount preparations of young wildtype and HCAR2/GPR109A knockout mouse retina (Figure 8). Very few leukocytes (FITC-ConA positive cells) were detected in wildtype retinas. In contrast, numerous adherent leukocytes were detected in HCAR2/GPR109A knockout flatmounts as supported by quantitation of FITC-ConA-positive cells in flatmount images (Fig. 8B). Glial cell activation and macrophage infiltration are also crucial to the retinal inflammatory response. We have already demonstrated upregulated GFAP immunolabeling in knockout mouse retinas (Figs. 4A-C). To evaluate macrophage/microglia numbers in wildtype and HCAR2/GPR109A knockout mouse retinas, retinal flatmounts were immunolabeled with Iba-1 (ionized calcium-binding adapter molecule 1 also known as allograft inflammatory factor 1, AIF-1; Fig. 8C). Significantly more Iba-1-positive (red-fluorescence) cells were detected in HCAR2/GPR109A knockout mouse retinas. This was confirmed by flow cytometric identification of cells that were CD11b and F4/80 positive (upper right quadrant) in single cell suspensions prepared from retinal homogenates. A representative flow tracing is provided in Fig. 8D. These data indicate an increase in immune cell activation and migration into HCAR2/GPR109A knockout mouse retina under basal conditions (in the absence of exogenous inflammatory stimuli), a phenomenon supportive of the functional and morphological phenotype observed and highly congruent with gene microarray and protein interactome analyses.

**Figure 8.**
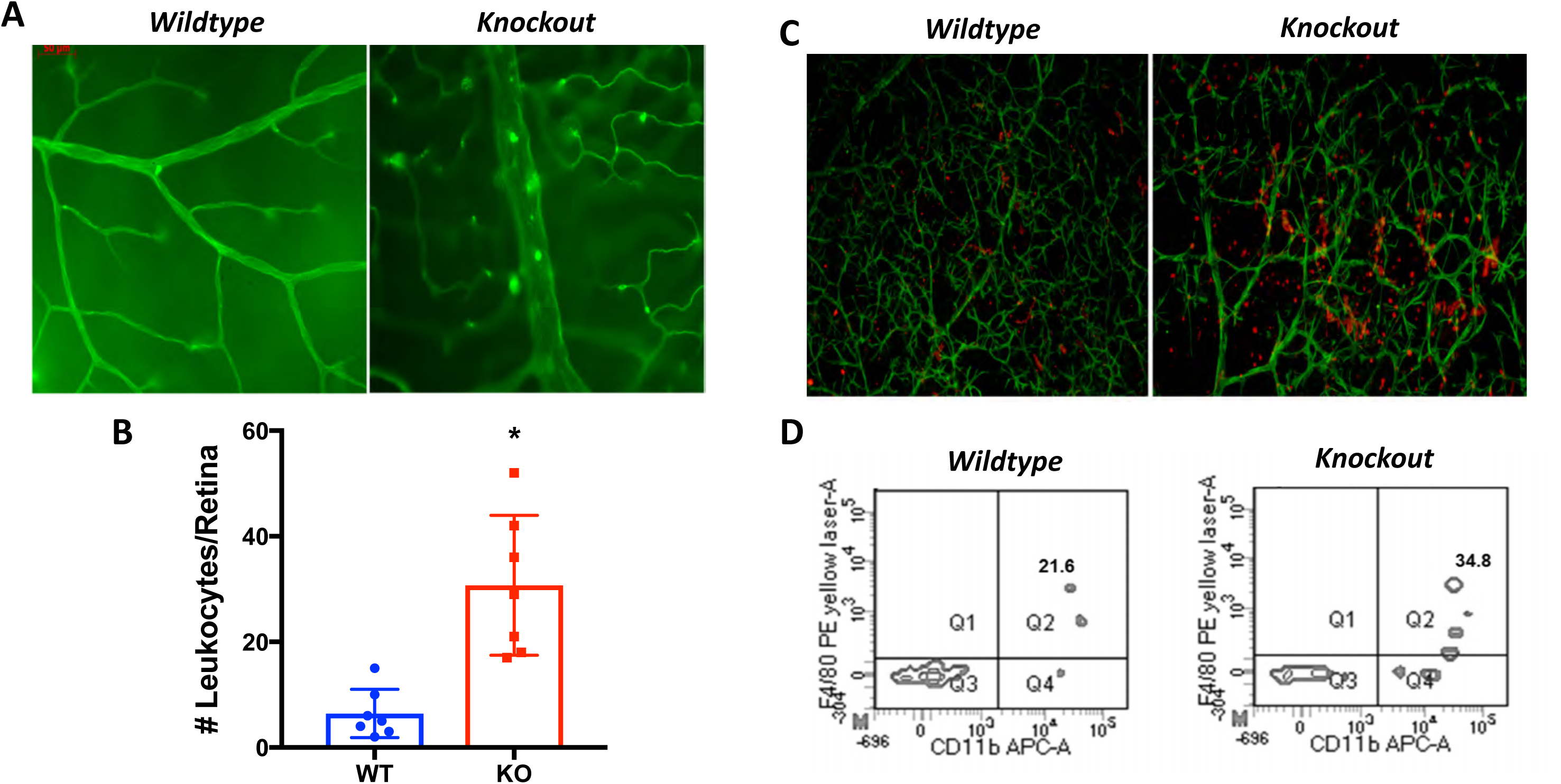
Increased activation and infiltration of inflammatory immune cells is common in HCAR2/GPR109A knockout mouse retina. (*A*) Representative retinal flatmounts prepared following leukostasis assay in 8-week old mice. Few FITC-Concanavalin A (FITC-ConA)-positive cells were detected in flatmount preparations of wildtype (WT) mouse retinas compared to HCAR2/GPR109A knockout (KO) mouse retinas. (*B*) Leukocytes that were adhered to the retinal vasculature were quantified in 6-9 flat mount preprations of each experimental condition by a blinded observer. Significantly more adherent leukocytes in HCAR2/GPR109A KO retina compared to WT retinas (*p<0.01). *(C)* Glial cell activation and macrophage infiltration was also monitored by immunolabeling for GFAP (green) and IBA-1 (red), a microglia/macrophage-specific marker. *(D)* To quantify immune cell populations in HCAR/GPR109A KO and WT mouse retinas, additional mice were perfused with PBS and retinas were collected and homogenized into single-cell suspensions and subjected to flow cytometry cell sorting analyses. A representative image of a CD45+ gated cells shows increase in CD11b+ and F4/80+ cells, a population of highly inflammatory macrophages in KO retinas.

### Therapeutic targeting of HCAR2/GPR109A suppresses immune cell activation and migration in retina

Since absence of HCAR2/GPR109A expression is associated with an increase in unregulated inflammatory responses, it seems that when present, HCAR2/GPR109A or signaling mediated thereby imposes important regulatory control. Thus, targeting the receptor therapeutically in pro-inflammatory conditions should be an effective means of limiting inflammation in retina. To validate this hypothesis, we examined whether activation of the receptor using beta-hydroxybutyrate (BHB), a known HCAR2/GPR109A agonist, modified the recruitment of pro-inflammatory cells to the retina in the lipopolysaccharide (LPS)-induced uveitis model, a model used widely to represent acute retinal inflammation. Flow cytometric analyses of single cell suspensions prepared from retinal homogenates derived from wildtype and HCAR2/GPR109A knockout mice treated with PBS (control) or LPS in the presence or absence of BHB, revealed a significantly high number of adherent leukocytes in LPS-treated mouse retinas, both wildtype and HCAR2/GPR109A knockout (Fig. 9A-B). Inflammation/edema was so robust in LPS-treated knockout animals that upon visualization by fluorescence microscopy background fluorescence was especially high. Many FITC-ConA positive leukocytes could be detected however it was obvious that the impact of LPS atop the already inflammatory environment likely to exist in normal knockout mouse retinas produced exacerbated damage as reflected by significant leakage of fluorescence dye into the retina. As such, images in Fig. 9A are presented in monochrome format to better highlight the leukocytes, now white in color. BHB treatment was effective in reducing leukocyte numbers in the retinas of wildtype but not HCAR2/GPR109A knockout mice treated with LPS. This is corroborated by flow cytometric analyses in which CD45+ cells were captured and gated again based upon Cd11b and Gr-1 positivity such that cells and corresponding values in the upper right quadrant of each tracing indicate the number of CD45+ positive cells that were specifically of a highly inflammatory phenotype in each retina (e.g., macrophages, neutrophils). A representative FACS tracing is providing in Fig. 9B.

**Figure 9.**
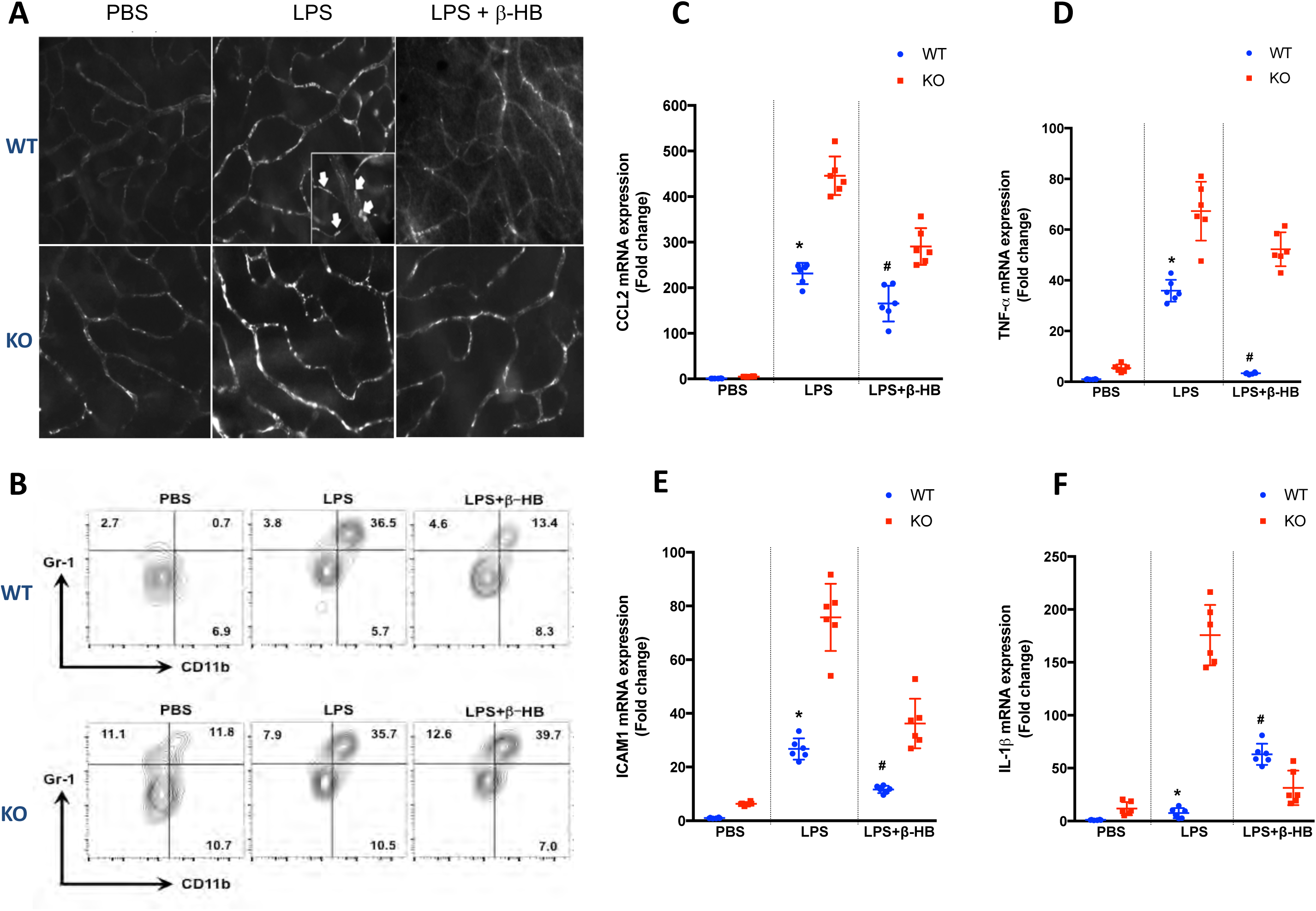
HCAR2/GPR109A expression and activation maintains an immunoinhibitory retinal environment. Wildtype and HCAR2/GPR109A knockout mice were treated intraperitoneally with PBS or with LPS conjunction with PBS or with the HCAR2/GPR109A agonist beta-hydroxybutyrate (BHB, 300 mg/kg) and the presence of retained leukocytes within the retinal vasculature was monitored via leukostasis assay using FITC-concanavalin A (FITC-ConA). Representative images of retinal flatmounts prepared from these mice are provided in *A.* (B) Retinal homogenates were additionally prepared from mice treated as described above and analyzed by flow cytometry to detect CD45+ cells that were additionally Cd11b and Gr-1 positive (upper right quadrant of each panel). (*C-F*) qPCR analyses of inflammatory molecule expression in mRNA samples obtained from mice treated as described above corroborated the increase in inflammation in LPS-treated wildtype and HCAR2/GPR109A knockout mice.

## Discussion

Collectively our results implicate HCAR2/GPR109A as an essential regulator of retinal immunity and tolerance under normal conditions and when the retina is faced with exogenous pathogens that illicit pro-inflammatory responses. These data are congruent with our prior published *in vitro* studies and studies by others demonstrating an anti-inflammatory role for this receptor in retina and other tissues. In the absence of HCAR2/GPR109A expression, functional deficits were detected in multiple retinal components (a-, b- and c-wave) early on and worsened progressively with age. *In vivo* imaging and histological analyses of retinas of mice at comparable ages revealed widespread disruption of neuro-retinal and vascular morphology that explained the physiologic basis of these deficits. Because prior studies have demonstrated robust expression of HCAR2/GPR109A in the RPE, we looked in depth at the morphology of this important cellular layer. ZO-1 immunolabeling revealed numerous areas of disruption in the typical cobblestone appearing phenotype were noted in flatmount preparations of HCAR2/GPR109A knockout RPE. ZO-1 expression was also reduced at the RNA level. These data support our histological observations of intact retinal cryosections in which clear disruptions in the RPE/choroidal interface were readily apparent. Because the model examined is a global knockout of the receptor, we wondered whether breakdown of the BRB, inflammation and degeneration of the retina is a direct result of local events or stem indirectly from changes occurring systemically. However, follow-up studies conducted using primary RPE cells isolated from wildtype and HCAR2/GPR109A knockout mouse eyes that were subjected to extended culture conditions to facilitate cell re-polarization/differentiation implicate changes inherent to HCAR2/GPR109A knockout RPE. HCAR2/GPR109A knockout primary RPE failed to form the proper cell-cell junctional complexes necessary to maintain barrier integrity and limit the extravasation of FITC-Dextran dye. This confirms that knockout RPE in general have a decreased ability to form consistent and appropriate tight junctions given that these cells are isolated from very young animals (18-21 days of age). Because we now know that in addition to RPE, HCAR2/GPR109A is expressed also by retinal microglia and endothelial cells (3,8,10), we also evaluated the integrity of retinal endothelial cells in trypsin digested preparations of wildtype and HCAR2/GPR109A knockout mouse retina. As with RPE, a number of histologic abnormalities indicative of endothelial cell dysfunction (e.g., acellular and/or degenerate capillaries) were evident. Collectively, these data suggest that outer- and inner-retinal barrier dysfunction occurs in association with absence of HCAR2/GPR109A expression.

Our present *in vivo* studies support strongly prior in vitro findings of an anti-inflammatory role for HCAR2/GPR109A. In effort to identify the principal molecular mechanisms responsible for this function in retina, gene microarray and protein interactome studies were performed using tissues isolated from wildtype and HCAR2/GPR109A knockout mouse eyes. Of the many genes that were differentially regulated it is interesting that those relevant to immune cell activation and homing to sites of inflammation stood out as some of the most significant. Leukostasis and flow cytometric analyses corroborated the presence of significantly more pro-inflammatory immune cells in HCAR2/GPR109A knockout mouse retinas under basal conditions. Further, in the LPS model of acute retinal inflammation, we found that treatment with beta-hydroxybutyrate (BHB), a well-known HCAR2/GPR109A receptor agonist, effectively suppressed the infiltration of these cells into retinal tissue. Because BHB was effective in wildtype mice but not in HCAR2/GPR109A mice, we can reliably state that the effects were due to HCAR2/GPR109A-mediated signaling. Others reported recently on the efficacy of BHB therapy against retinal dysfunction in experimental diabetes in rodents (25), a chronic inflammatory condition and concluded that the effects too stemmed from HCAR2/GPR109A activation. These findings are in line with those of this study however, it is important that they be confirmed given that only wild type animals (no HCAR2/GPR109A knockout or knockdown) were used to differentiate definitively between receptor-dependent and receptor-independent effects.

G-protein coupled receptors (GPCRs) elicit a myriad of signaling responses, some good and some bad. Importantly, they do so by interacting with molecules, circulating within the extracellular environment. Hence, their robust and successful pharmacologic targeting of GPCRs to treat a number of systemic diseases. The present work demonstrates for the first time the functional relevance of HCAR2/GPR109A expression in retina. The robust expression of HCAR2/GPR109A in barrier and immune cells puts HCAR2/GPR109A forth as a viable potential clinical target for the prevention and treatment of retinal disease. Regardless of the limitations mentioned above, our current study coupled with those of others yield promise that a means can be developed to therapeutically impact inflammation in retina other than intravitreal injection, the current gold standard for administering drugs to retina. In the present study, we administered BHB intraperitoneally. However, others and ourselves (unpublished result) have also demonstrated the ability to raise BHB levels high enough to impact HCAR2/GPR109A activation via dietary modifications such as ketogenic diet, caloric restriction or intermittent fasting (9,27,28). The potential clinical significance of this is enormous given (a) that dysregulated immune responses and related inflammation is a major pathogenic component of degenerative retinal disease due to aging, diabetes, glaucoma, sickle cell, etc. and, (b) the known risks associated with the invasive delivery of drugs intravitreally (e.g., injury, infection, scarring).

## Abbreviations

BHB: beta-hydroxybutyrate
ConA: concanavalin A
FA: fluorescein angiography
GPCR: G-protein coupled receptor
Gr-1: Ly-6G/Ly-6C
IbA-1: ionized calcium-binding adapter molecule 1 also known as AIF-1, allograft inflammatory factor 1
LPS: lipopolysaccharide
OCT: spectral domain optical coherence tomography
RPE: retinal pigmented epithelium
ZO-1: zonula occludens -1.

## References

1. Al-Shabrawey M, Rojas M, Sanders T, Behzadian A, El-Remessy A, Bartoli M, Parpia AK, Liou G, Caldwell RB. Role of NADPH oxidase in retinal vascular inflammation. Invest Ophthalmol Vis Sci 49: 3239–44, 2008.

2. Catorce MN, Gevorkian G. LPS-induced Murine Neuroinflammation Model: Main Features and Suitability for Pre-clinical Assessment of Nutraceuticals. Curr Neuropharmacol 14: 155–64, 2016.

3. Chen H, Assmann JC, Krenz A, Rahman M, Grimm M, Karsten CM, Kohl J, Offermanns S, Wettschureck N, Schwaninger M. Hydroxycarboxylic acid receptor 2 mediates dimethyl fumarate’s protective effect in EAE. J Clin Invest 124: 2188–92, 2014.

4. Cresci GA, Thangaraju M, Mellinger JD, Liu K, Ganapathy V. Colonic gene expression in conventional and germ-free mice with a focus on the butyrate receptor GPR109A and the butyrate transporter SLC5A8. J Gastrointest Surg 14: 449–61, 2010.

5. Elangovan S, Pathania R, Ramachandran S, Ananth S, Padia RN, Lan L, Singh N, Martin PM, Hawthorn L, Prasad PD, Ganapathy V, Thangaraju M. The niacin/butyrate receptor GPR109A suppresses mammary tumorigenesis by inhibiting cell survival. Cancer Res 74: 1166–78, 2014.

6. Gambhir D, Ananth S, Veeranan-Karmegam R, Elangovan S, Hester S, Jennings E, Offermanns S, Nussbaum JJ, Smith SB, Thangaraju M, Ganapathy V, Martin PM. GPR109A as an anti-inflammatory receptor in retinal pigment epithelial cells and its relevance to diabetic retinopathy. Invest Ophthalmol Vis Sci 53: 2208–17, 2012.

7. Harun-Or-Rashid M, Inman DM. Reduced AMPK activation and increased HCAR activation drive anti-inflammatory response and neuroprotection in glaucoma. J Neuroinflammation 15: 313, 2018.

8. Hughes-Large JM, Pang DK, Robson DL, Chan P, Toma J, Borradaile NM. Niacin receptor activation improves human microvascular endothelial cell angiogenic function during lipotoxicity. Atherosclerosis 237: 696–704, 2014.

9. Izuta Y, Imada T, Hisamura R, Oonishi E, Nakamura S, Inagaki E, Ito M, Soga T, Tsubota K. Ketone body 3-hydroxybutyrate mimics calorie restriction via the Nrf2 activator, fumarate, in the retina. Aging Cell 17, 2018.

10. Jiang D, Ryals RC, Huang SJ, Weller KK, Titus HE, Robb BM, Saad FW, Salam RA, Hammad H, Yang P, Marks DL, Pennesi ME. Monomethyl Fumarate Protects the Retina From Light-Induced Retinopathy. Invest Ophthalmol Vis Sci 60: 1275–1285, 2019.

11. Martin PM, Ananth S, Cresci G, Roon P, Smith S, Ganapathy V. Expression and localization of GPR109A (PUMA-G/HM74A) mRNA and protein in mammalian retinal pigment epithelium. Mol Vis 15: 362–72, 2009.

12. Martin PM, Gambhir D, Promsote W, Ganapathy V, Moore-Hill D. Expression of the niacin receptor GPR109A in retina: more than meets the eye. Clinical Experimental Pharmacology, 2013.

13. Parodi B, Rossi S, Morando S, Cordano C, Bragoni A, Motta C, Usai C, Wipke BT, Scannevin RH, Mancardi GL, Centonze D, Kerlero de Rosbo N, Uccelli A. Fumarates modulate microglia activation through a novel HCAR2 signaling pathway and rescue synaptic dysregulation in inflamed CNS. Acta Neuropathol 130: 279–95, 2015.

14. Pike NB, Wise A. Identification of a nicotinic acid receptor: is this the molecular target for the oldest lipid-lowering drug? Curr Opin Investig Drugs 5: 271–5, 2004.

15. Promsote W, Powell FL, Veean S, Thounaojam M, Markand S, Saul A, Gutsaeva D, Bartoli M, Smith SB, Ganapathy V, Martin PM. Oral Monomethyl Fumarate Therapy Ameliorates Retinopathy in a Humanized Mouse Model of Sickle Cell Disease. Antioxid Redox Signal 25: 921–935, 2016.

16. Promsote W, Veeranan-Karmegam R, Ananth S, Shen D, Chan CC, Lambert NA, Ganapathy V, Martin PM. L-2-oxothiazolidine-4-carboxylic acid attenuates oxidative stress and inflammation in retinal pigment epithelium. Mol Vis 20: 73–88, 2014.

17. Roy S, Kim D, Hernandez C, Simo R, Roy S. Beneficial effects of fenofibric acid on overexpression of extracellular matrix components, COX-2, and impairment of endothelial permeability associated with diabetic retinopathy. Exp Eye Res 140: 124–129, 2015.

18. Saul AB, Humphrey AL. Spatial and temporal response properties of lagged and nonlagged cells in cat lateral geniculate nucleus. J Neurophysiol 64: 206–24, 1990.

19. Schaub A, Futterer A, Pfeffer K. PUMA-G, an IFN-gamma-inducible gene in macrophages is a novel member of the seven transmembrane spanning receptor superfamily. Eur J Immunol 31: 3714–25, 2001.

20. Singh N, Gurav A, Sivaprakasam S, Brady E, Padia R, Shi H, Thangaraju M, Prasad PD, Manicassamy S, Munn DH, Lee JR, Offermanns S, Ganapathy V. Activation of Gpr109a, receptor for niacin and the commensal metabolite butyrate, suppresses colonic inflammation and carcinogenesis. Immunity 40: 128–39, 2014.

21. Soga T, Kamohara M, Takasaki J, Matsumoto S, Saito T, Ohishi T, Hiyama H, Matsuo A, Matsushime H, Furuichi K. Molecular identification of nicotinic acid receptor. Biochem Biophys Res Commun 303: 364–9, 2003.

22. Taggart AK, Kero J, Gan X, Cai TQ, Cheng K, Ippolito M, Ren N, Kaplan R, Wu K, Wu TJ, Jin L, Liaw C, Chen R, Richman J, Connolly D, Offermanns S, Wright SD, Waters MG. (D)-beta-Hydroxybutyrate inhibits adipocyte lipolysis via the nicotinic acid receptor PUMA-G. J Biol Chem 280: 26649–52, 2005.

23. Tang H, Lu JY, Zheng X, Yang Y, Reagan JD. The psoriasis drug monomethylfumarate is a potent nicotinic acid receptor agonist. Biochem Biophys Res Commun 375: 562–5, 2008.

24. Thangaraju M, Cresci GA, Liu K, Ananth S, Gnanaprakasam JP, Browning DD, Mellinger JD, Smith SB, Digby GJ, Lambert NA, Prasad PD, Ganapathy V. GPR109A is a G-protein-coupled receptor for the bacterial fermentation product butyrate and functions as a tumor suppressor in colon. Cancer Res 69: 2826–32, 2009.

25. Trotta MC, Maisto R, Guida F, Boccella S, Luongo L, Balta C, D’Amico G, Herman H, Hermenean A, Bucolo C, D’Amico M. The activation of retinal HCA2 receptors by systemic beta-hydroxybutyrate inhibits diabetic retinal damage through reduction of endoplasmic reticulum stress and the NLRP3 inflammasome. PLoS One 14: e0211005, 2019.

26. Tunaru S, Kero J, Schaub A, Wufka C, Blaukat A, Pfeffer K, Offermanns S. PUMA-G and HM74 are receptors for nicotinic acid and mediate its anti-lipolytic effect. Nat Med 9: 352–5, 2003.

27. Youm YH, Nguyen KY, Grant RW, Goldberg EL, Bodogai M, Kim D, D’Agostino D, Planavsky N, Lupfer C, Kanneganti TD, Kang S, Horvath TL, Fahmy TM, Crawford PA, Biragyn A, Alnemri E, Dixit VD. The ketone metabolite beta-hydroxybutyrate blocks NLRP3 inflammasome-mediated inflammatory disease. Nat Med 21: 263–9, 2015.

28. Zarnowski T, Choragiewicz TJ, Schuettauf F, Zrenner E, Rejdak R, Gasior M, Zarnowska I, Thaler S. Ketogenic diet attenuates NMDA-induced damage to rat’s retinal ganglion cells in an age-dependent manner. Ophthalmic Res 53: 162–7, 2015.

